# Expectation for sweet taste changes peripheral glucose metabolism via basolateral amygdala

**DOI:** 10.1101/2022.02.03.479066

**Authors:** Izumi Yamamoto, Toya Yonekura, Taiga Ishimoto, Shu-Cheng Xu, Norifumi Iijima, Kazuhiro Kimura, Sabrina Diano, Chitoku Toda

## Abstract

Anticipatory physiological responses to food were first reported by Ivan Pavlov a century ago but the associated neural mechanism is still ill-defined. Here, we identified two types of neurons in the basolateral amygdala (BLA), which are activated by sweetener (saccharin) or water after sucrose conditioning, representing expected sweet taste and unmet expectation, respectively. Saccharin-induced met-expectation of sweet taste enhances, while H_2_O-induced unmet-expectation deteriorates, glucose metabolism in peripheral tissues. Deletion of saccharin-responsive neurons in BLA impaired saccharin-induced increase in insulin sensitivity. Deletion of H_2_O-responsive neurons in BLA improved glucose intolerance by unmet-expectation. Saccharin- and H_2_O-responsive neurons had different gene expressions. Our data suggest that the gap between the expected incoming sugar and sweet taste is evaluated by distinct BLA neurons to control peripheral glucose metabolism.

**One-Sentence Summary:** Neurons in the basolateral amygdala control blood glucose levels by comparing anticipated sugar intake and sweet taste

## Introduction

According to Pavlov’s dog experiments documented in the early 20th century, animals predict the start of a meal and show physiological responses. Pavlov showed that when food-independent sound cues are associated with food, a representation of the cue increases the secretion of saliva, gastric acid, pancreatic enzymes, etc (*1*, *2*). Similar effects also occur when food is perceived through sight, smell, and taste (*3*). These reactions are called cephalic responses and enable efficient digestion, absorption, and nutritional regulation before the start of food intake (*4*). Cephalic responses play an important role in the brain’s regulation of systemic energy metabolism (*5*).

Peripheral hormones and nutrients, which reflect the amount of energy availability in the body, are known to affect the activity of neurons in order to regulate food intake and energy utilization in peripheral tissues (*6*). Among the peripheral hormones, insulin plays a particularly important role in glucose metabolism. When sugar is expected to be ingested, the cephalic phase insulin release (CPIR) increases to prepare for the control of blood glucose levels before food intake (*7*–*9*). Studies in humans suggest that taste is deeply involved in the regulation of blood glucose levels by cephalic responses.

Pseudo-feeding (chew and spit) enhances glucose metabolism by a pathway that does not involve an increase in insulin secretion (*10*), and oral sensory stimulation is accompanied by an increase in insulin secretion to enhance glucose metabolism (*9*, *11*, *12*). The effect of the cephalic responses on glucose metabolism is very fast, suggesting that the parasympathetic nervous system plays an important role in the regulation of CPIR (*7*, *8*, *13*). It has also been suggested that the hypothalamus and vagus nerve are important as the neural mechanism of CPIR (*8*), but the whole picture of the neuronal mechanism that controls cephalic responses is not clear.

Regulation of systemic glucose metabolism by cephalic responses is important to prevent acute hyperglycemia after a meal. Postprandial hyperglycemia and chronic hyperglycemia are risk factors for diabetic complications (*14*, *15*). It has been suggested that not only abnormal blood glucose levels, but also psychological factors such as depression and anxiety are risk factors for the development of type II diabetes (*16*–*19*). Daily psychological emotions can also change the blood glucose level of diabetic patients, i.e., positive emotions lower blood glucose levels, and negative emotions increase blood glucose levels (*20*–*22*). However, the neural mechanism by which psychological factors affect blood glucose levels and cephalic responses have not been clarified yet.

## Materials and Methods

### Animals

All animal experiments were approved by the Animal Care and Use Committee of Hokkaido University and were performed according to the institutional guidelines. Arc-Cre/ER^T2^ mice (Strain #021881, Jackson laboratory, Bar Harbor, ME, USA) and Ai14 mice (Strain #007914, Jackson laboratory) were crossed to obtain Cre/ERT2;Ai14 mice. Female Arc-Cre/ER^T2^;Ai14 mice (9-50 weeks of age) were used. All mice were kept at 22 ± 4 °C under a 12/12-h light/dark cycle (light phase: 7:00– 19:00) and given ad libitum food and water access. Mice were handled using the tunnel method to reduce stress and anxiety in the mice during experiments.

### Sucrose conditioning

For the conditioning to conduct open field test, a nontransparent square box (38 × 38 cm^2^) with a yellow mark at one of the corners was used. A mouse was placed on the opposite side of the yellow mark and the mouse was allowed to explore the arena for 3 min. 1 mL of 15% sucrose solution (15% w/v, Fujifilm Wako Pure Chemical Industries, Osaka, Japan) was set under the yellow mark and kept for 5 min. All the mice found the sucrose solution within 4 min in the first trial. Training sessions were performed once a day for 5-8 consecutive days.

For the conditioning to measure glucose metabolism and TRAP, a mouse was placed in a normal plastic cage that was used for the home cage (30.6 × 19.6 cm^2^, MBS75JRH, Allentown Caging, NJ) without bedding. The mouse was allowed to explore the cage for 3 min and a water bottle with 15% sucrose solution (15% w/v, Fujifilm Wako Pure Chemical Industries) was inserted into the cage for 5 min. All the mice found the sucrose solution within 4 min in the first trial. Training sessions were performed once a day for 5-8 consecutive days. On the test day, the mouse was placed in the cage without bedding and allowed to explore for 3 min. A water bottle with 0.015% saccharin solution (Fujifilm Wako Pure Chemical Industries) or H_2_O was inserted, and a glucose tolerance test was started.

### Open field test

On the test day, sucrose solution or H_2_O was set under the yellow mark, and the mouse was placed on the opposite side of the yellow mark to measure the latency to drink solutions first time. The behavior of the mouse was video-recorded for 5 min and analyzed using ImageJ [15] and MouBeAT [18]. The weight of the solution was measured to assess the solution intake.

## Targeted recombination in active populations (TRAP)

After 8 days of conditioning with sucrose, female Arg3.1-Cre/ER^T2^;Ai14 mice were received 4-hydroxy tamoxifen (4-OHT) (10 mg/kg; Sigma-Aldrich, St. Louis, MO, USA) intraperitoneal (i.p.) injection and were kept in the conditioning cage for 3 min. A water bottle with 0.015% saccharin solution or H_2_O was inserted. Mice were returned to their home cage 60 min after drinking one of the solutions. To ensure adequate expression of tdTomato proteins, experiments were carried out at least 7 days after the 4-OHT injection.

### Glucose tolerance test and insulin tolerance test

For the glucose tolerance test, animals were fasted overnight and injected with glucose (2 g/kg; i.p.). Blood glucose levels were measured with a handheld glucose meter (Nipro Free Style, Nipro, Osaka, Japan) at 0, 15, 30, 60 and 120 min after glucose injection. The insulin tolerance test was performed on ad libitum-fed mice. The mice were injected with 0.75 U/kg insulin (Novo Nordisk, Bagsværd, Denmark). Blood glucose was measured at 0, 15, 30, 60 and 120 min after insulin injection. Some mice (saccharin group in Fig. 3A) received a glucose injection and the experiment was stopped when the blood glucose level became under 30 mg/dl.

### Serum C-peptide

Blood (40 μL) from the tails was taken at 0 and 15 min of the GTT. The serums were collected after centrifuging for 5 min at 3000 × *g* and maintained at−80°C until C-peptide was measured. The C-peptide concentration was measured with a Mouse C-peptide ELISA Kit (M1304, Morinaga Institute of Biological Science, Inc., Tokyo, Japan) and all procedures were performed according to the protocols provided in the kit.

### Brain sectioning

Mice were sacrificed using CO_2_ asphyxiation and perfused with heparinized saline. Brain samples were harvested and incubated in 4% paraformaldehyde (PFA) overnight. Brain sections (50 μm) were collected, rinsed with 0.1 M phosphate buffer, and mounted on glass slides with Vectashield (Vector Laboratories, Burlingame, CA, USA).

### Stereotaxic surgeries and adeno-associated virus (AAV) injection

Arc-Cre/ER^T2^ mice (12–20-week-old) were anesthetized with a mixture of ketamine (100 mg/kg) and xylazine (10 mg/kg) and positioned on a stereotaxic instrument (Narishige, Tokyo, Japan). Mice were injected in both sides of the basolateral amygdala (BLA) with ~0.5 μL AAV2-Flex-taCasp3-Tevp (UNC vector core, NC) using the following coordinates: Anterior-Posterior: −1.83 mm, Lateral: ±3.3 mm, Dorsal-Ventral: −5.0 mm. Open wounds were sutured after viral injection. A 7–14-day recovery period was allowed before experiments were started. Mice received a 4-OHT (10 mg/kg) injection (i.p.) as described above to induce Cre recombination. To ensure adequate expression of proteins, experiments were carried out at least 7 days after the 4-OHT injection.

### Cell sorting for RNA sequencing

TRAP-labeled saccharin- or H_2_O-responsive neurons were collected by sorting tdTomato-positive neurons from the BLA of female Arc-Cre/ER^T2^;Ai14 mice. We used a modified protocol for manual sorting [19]. Mice were sacrificed using CO_2_ asphyxiation, and the brain was placed in ice-cold cutting solution (220 mM sucrose, 2.5 mM KCl, 6 mM MgCl_2_, 1 mM CaCl_2_, 1.25 mM NaH_2_PO_4_, 10 mM glucose, 26 mM NaHCO_3_, bubbled thoroughly with 95% O_2_/5% CO_2_). A coronal brain slice (500 μm) containing the BLA was obtained using a vibratome (PELCO easiSlicer, Redding, CA, USA). The BLA was dissected and placed in a tube containing filtered artificial cerebrospinal fluid (ACSF; 105 mM NaCl, 2.5 mM KCl, 2 mM CaCl_2_, 1.3 mM MgSO_4_, 1.23 mM NaH_2_PO_4_, 24 mM NaHCO_3_, 20 mM HEPES, 2.5 mM glucose, 100 nM TTX, 20 μM DQNX, 50 μM AP-V, pH 7.4, bubbled thoroughly with 95% O_2_/5% CO_2_) with papain (0.3 U/mL), DNase (0.075 μg/mL) and BSA (3.75 μg/mL). The tube was incubated on a shaker (34 °C, 75 rpm) for 15 min. After incubation, the tissue was washed three times with papain-free filtered ACSF supplemented with FBS (1%). Subsequently, 1 mL of ACSF with FBS (1%) was added, and the samples were triturated successively with 600, 300 and 150 μm fire-polished Pasteur pipettes (10 times each). The tube was centrifuged (120 × *g*, 5 min, room temperature), and the supernatant was removed. The cell pellet was resuspended in 10 mL of filtered ACSF containing FBS (1 %) and transferred to a 100 mm collagen type I-coated petri dish. The cells were allowed to settle onto the floor of the dish for 10 min, and tdTomato-positive neurons were sorted separately under the fluorescence microscope using a cell aspirator attached to a micropipette (diameter: 30–50 μm). The neurons were transferred to a clean 35 mm collagen type I-coated petri dish containing filtered ACSF with FBS (1%). This sorting process was repeated one more time, and cells were transferred to a 35 mm collagen type I-coated petri dish containing filtered PBS (bubbled thoroughly with 95% O_2_/5% CO_2_). The sorted neurons were transferred to a PCR tube (one neuron per tube) containing 1 μL of 10× reaction buffer (SMART-Seq v4 Ultra Low Input RNA Kit for Sequencing, Cat. No. 634888, Takara Bio, Kusatsu, Japan). Nuclease-free water was added to bring the total volume to 11.5 μL, and the mixture was stored at −80 °C until library preparation.

### RNA sequencing

RNA sequence library preparation, sequencing, and mapping of gene expression were performed by DNAFORM (Yokohama, Kanagawa, Japan). Double-stranded cDNA libraries (RNA-seq libraries) were prepared using SMART-Seq v4 Ultra Low Input RNA Kit for Sequencing (Cat. No. 634888, Takara Bio) according to the manufacturer’s instructions. RNA-seq libraries were sequenced using paired-end reads (50-nt read 1 and 25-nt read 2) on a NextSeq 500 instrument (Illumina, San Diego, CA, USA). Obtained reads were mapped to the mouse GRCm38.p6 genome using STAR (version 2.7.3a) [20]. Reads on annotated genes were counted using featureCounts (version 1.6.4) [21].

FPKM values were calculated from mapped reads by normalizing to total counts and transcript. Prism 8 software (GraphPad) was used to generate heatmaps. Gene lists used for RNA sequencing analysis were obtained from previous reports [22,23].

### Statistical analysis

All data are presented as the mean ± SEM, and n represents the number of animals. Statistical differences were evaluated with Student’s unpaired *t*-test (for two-group comparisons) or two-way ANOVA. Sidak post-hoc tests (for multiple comparisons) were performed using Prism 9 software (GraphPad). Values of *P* < 0.05 were considered significant.

## Result

In this study, a sweet taste positive reinforcer (sucrose solution) was used to acquire a cue-sucrose association. Mice were placed in a box with yellow tape on the wall (an environmental cue) indicating the position of 15 % sucrose solution (Fig. 1A). After the conditioning with a cue-sucrose association, mice displayed a shorter latency time for the first lick of sucrose solution, a longer time staying near the sucrose solution, and higher sucrose intake compared with the mice before conditioning (Fig. 1B and C). To assess the effect of different sweet tastes, we used water as a non-flavored control (absence of an expected sweet taste positive reinforcer) (Fig. 1D). The latency time to drink H_2_O was similar to that of sucrose (Fig. 1E), but the mice stayed shorter and drank less when H_2_O was given compared with sucrose (Fig. 1F). Plasma corticosterone level was comparable between groups after saccharin or H_2_O drinking (Fig. 1G).

**Fig. 1.**
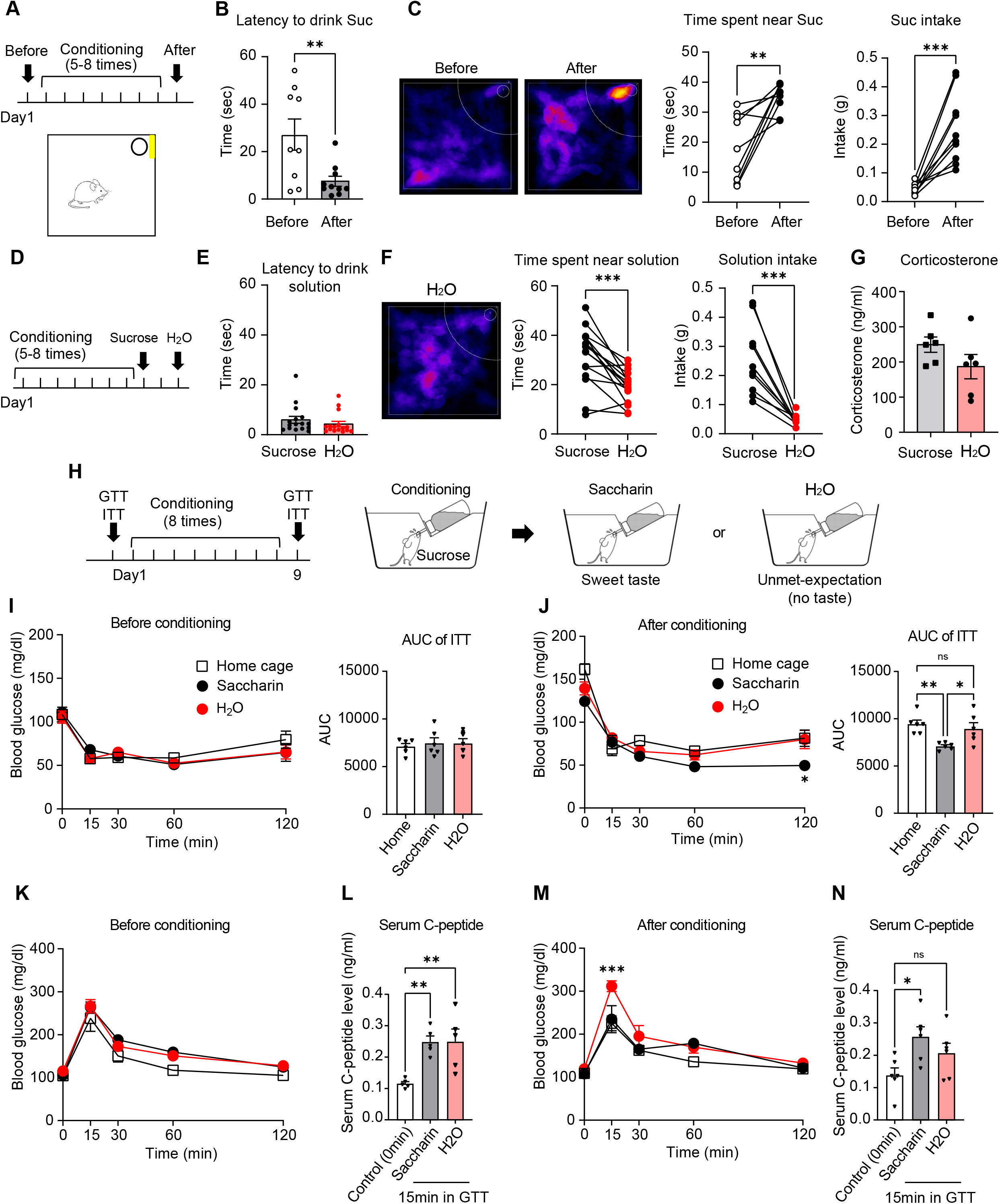
Met/unmet expectations for sweet taste influence systemic glucose metabolism. (A) Sucrose conditioning and open field test for sucrose drinking. (B) Latency to the first bout of sucrose consumption. (C) Representative image of an open field test, time spent near the sucrose solution, and the amount of sucrose intake. (D) Sucrose conditioning and the day to measure solution intake. (E-F) Open field test for sucrose and water intake. (G) Plasma corticosterone concentration after sucrose or H_2_O drinking. (H) Protocol of conditioning and GTT/ITT. (I) ITT without the sucrose conditioning and the area under the curve (AUC). Home cage: ITT was performed in the home cage. Saccharin and H_2_O: Mice were moved to the cage without bedding and saccharin solution or H_2_O was dropped in the cage. ITT was started 5min later. (J) ITT after 8 days of the sucrose conditioning and the AUC. Home cage: ITT was performed in the home cage. Mice would not expect a sucrose intake. Saccharin and H_2_O: Mice were moved to the cage without bedding and saccharin solution or H_2_O was dropped in the cage. ITT was started 5 min later. (K) GTT without the sucrose conditioning. (L) C-peptide concentration at 0 and 15 min of GTT in K. (M-N) GTT and C-peptide concentration after 8 days of the sucrose conditioning. Data are shown as mean ± SEM. \**P* < 0.05, \*\**P* < 0.01, \*\*\**P* < 0.001; two-tailed *t*-test in B, C, F; two-way ANOVA followed by Sidak multiple comparison test in I-N.

To check the effect of met/unmet-expectations of sweet taste on systemic glucose metabolism, insulin (ITT) and glucose (GTT) tolerance tests were performed. We trained a group of mice using an empty cage with no bedding as an environmental cue, and sucrose was delivered using a bottle. For GTT and ITT, saccharin, a zero-calorie sweetener, was used instead of sucrose as its calories may change blood glucose levels (Fig. 1H). We first performed GTT and ITT on unconditioned mice, using mice with an unpaired environmental cue (home cage) as a control group, and did not observe any differences in blood glucose levels among the three groups (Fig. 1I and K). In conditioned mice, the insulin sensitivity in the saccharin group was significantly enhanced at 120 mins after insulin injection as compared to H_2_O or control group (Fig. 1J). The area under the curve (AUC) of the saccharin group was also significantly decreased compared to other groups (Fig. 1J), suggesting that the Pavlovian conditioning increases insulin sensitivity to prepare for the expected sugar intake. Glucose tolerance was not different between saccharin and home cage groups, while mice which were given H_2_O had impaired glucose tolerance at 15 mins after glucose injection compared with the home cage group (Fig. 1M). Serum C-peptide level at 15min of GTT, which indicates insulin release, was significantly increased in the saccharin group, but not in the H_2_O group (Fig. 1M and N), suggesting that unmet-expectation suppresses insulin secretion from the pancreas in response to glucose injection.

To investigate the neuronal mechanisms of cephalic response to regulate blood glucose levels, we used the targeted recombination in active populations (TRAP) method (*23*). This method labels neurons that express activity-regulated cytoskeleton-associated protein (Arc/Arg3.1) gene, a member of the immediate-early gene family used as a marker of neuronal activity. Arc gene expression is induced by sensory experiences and also expressed in the amygdala (*24*, *25*). Arc-Cre/ER^T2^;Ai14 mice can express tdTomato for fluorescence visualization when 4-OHT, an active form of tamoxifen, was injected. Arc-Cre/ER^T2^;Ai14 mice were conditioned as described above. On the labeling day, the mice were injected with 4-OHT (i.p.) and then placed in the conditioning box, where they received saccharin solution or water (Fig. 2A). Unconditioned mice were also injected with 4-OHT (i.p.) as control. A significant increase in the number of tdTomato-positive basolateral amygdala (BLA) neurons was observed when the conditioned mice received saccharin solution or water compared to the control group, while there was no change between groups in the central amygdala (CeA) (Fig. 2B and C). The amygdala is where the brain processes positive and negative stimuli and plays a critical role in emotional behaviors (*26*, *27*). The BLA comprises various cell types and genetically diverse populations. To determine whether TRAP-labeled BLA neurons play a role in glucose homeostasis, we injected a Cre-dependent virus encoding pro-caspase 3 and TEV protease (AAV-Flex-taCasp3-Tevp) into the Arc-Cre/ER^T2^;Ai14 mice to selectively delete neurons responsive to H_2_O (ΔH_2_O-R) or saccharin (ΔSacc-R) (Fig. 3D). A significant reduction in tdTomato-positive basolateral amygdala (BLA) neurons was observed in ΔSacc-R and ΔH_2_O-R mice (Fig. 3E and F). Control animals were injected with a Cre-dependent GFP construct.

**Fig. 2.**
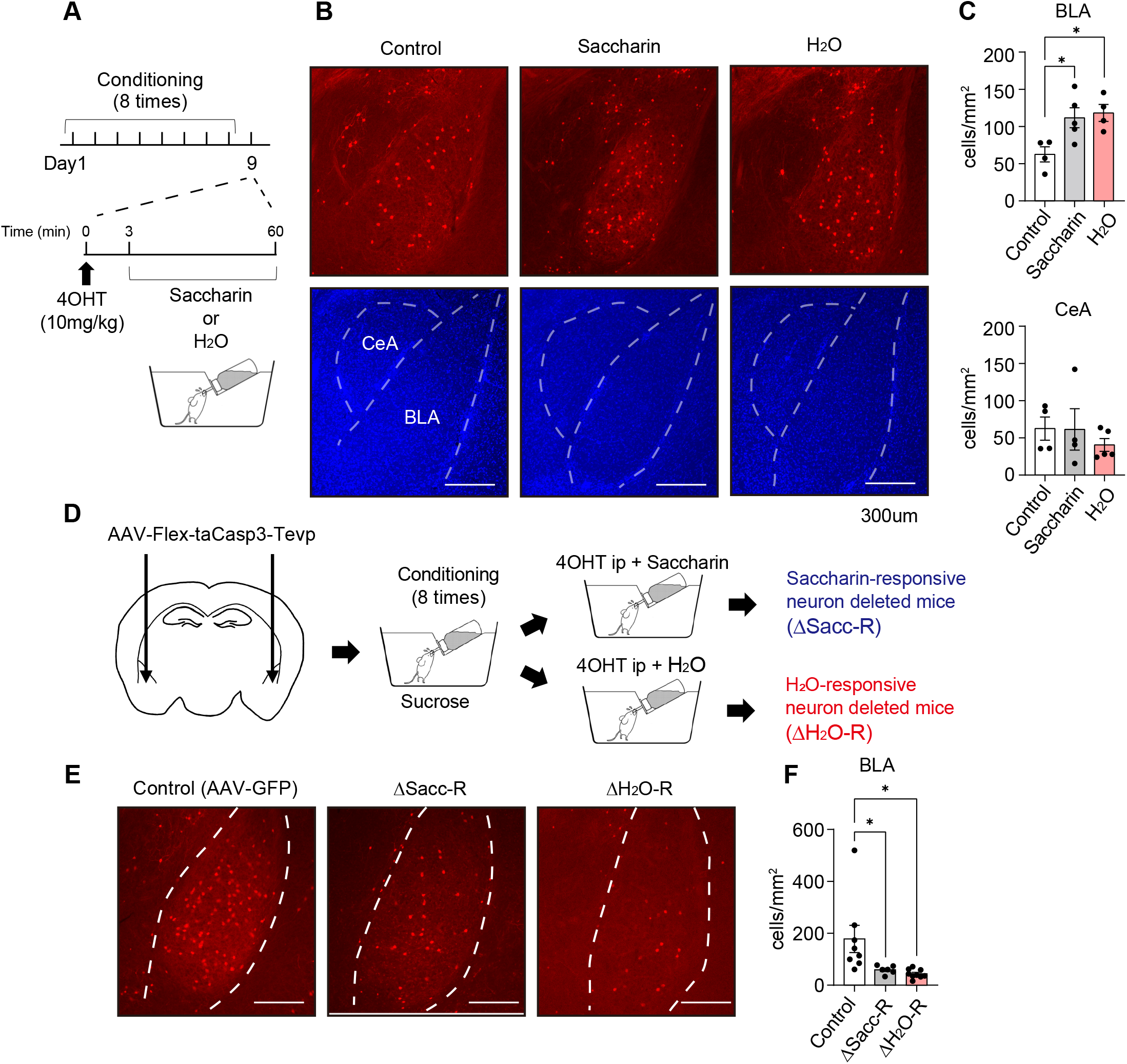
Met/unmet expectations for sweet taste activate neurons in the basolateral amygdala. (A) Protocol of the TRAP method. (B) Representative images of the basolateral amygdala (BLA) and central amygdala (CeA). (C) Quantified numbers of TRAPed neurons in BLA and CeA. (D) Schematic of AAV injection and strategy to delete H_2_O- or saccharin-responsive neurons in the BLA. (E-F) Representative image (E) and quantified numbers (F) of TRAPed neurons after the AAV-mediated cell deletion. Data are shown as mean ± SEM. \**P* < 0.05; two-way ANOVA followed by Sidak multiple comparison test.

**Fig. 3.**
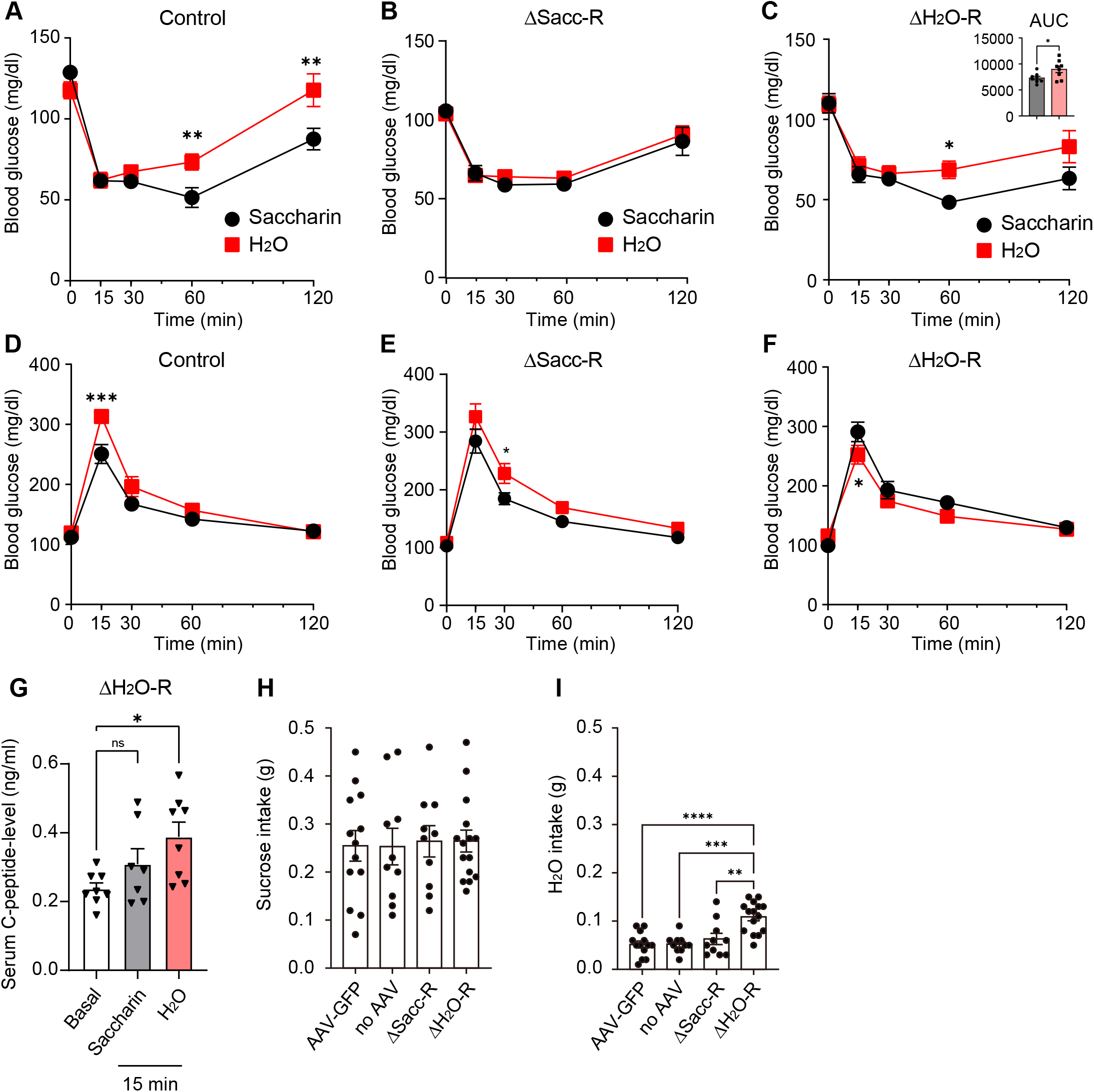
Ablation of TRAPed neurons suppresses the changes in glucose metabolism induced by the sweet taste. (A-C) ITT in control (A), H_2_O-responsive neuron deleted (B), and saccharin-responsive neuron deleted (C) mice after 7 days of sucrose conditioning. Saccharin and H_2_O: Mice were moved to the cage without bedding and saccharin solution or H_2_O was dropped in the cage. ITT was started 5 min later. (D-F) GTT in control (D), H_2_O-responsive neuron deleted (E), and saccharin-responsive neuron deleted (F) mice. (G) C-peptide concentration in H_2_O-responsive neuron deleted mice. (H-I) sucrose (H), and H_2_O (I) intake in the open field test after sucrose conditioning. Data are shown as mean ± SEM. \**P* < 0.05, \*\**P* < 0.01, \*\*\**P* < 0.001, \*\*\*\**P* < 0.0001; two-way ANOVA followed by Sidak multiple comparison test.

Then, we performed ITT and GTT after sucrose conditioning. In ITT, saccharin enhanced insulin sensitivity in control and ΔH_2_O-R mice (Fig. 3A and C), but not in ΔSacc-R mice (Fig. 3B), suggesting that saccharin-responsive neurons in the BLA regulate the Pavlovian conditioning-induced enhancement of insulin sensitivity. In GTT, H_2_O increased blood glucose levels at 15 min after the glucose injection in control and ΔSacc-R mice (Fig. 3D and E), however, the blood glucose level was significantly lower in the ΔH_2_O-R mice (Fig. 3F). Serum C-peptide level in ΔH_2_O-R mice was significantly increased by glucose injection even after inducing unmet-expectation by H_2_O drinking (Fig. 3G), which was not significant in Fig. 1M, suggesting that BLA neurons affect insulin secretion. These results suggest that these TRAP-labeled neurons in the BLA are critical for the met/unmet-expectation-related change in the regulation of peripheral glucose metabolism. The sucrose intake after the conditioning was not different after the deletion of TRAPed-BLA neurons (Fig. 3H). ΔH_2_O-R mice had increased H_2_O intake, suggesting that deletion of BLA neurons did not change the reward value of sucrose, while these neurons may affect the response to the unexpected change in taste.

To investigate the transcriptional profile of TRAPed-BLA neurons responding to saccharin and H_2_O, we performed an RNA-sequencing study using cells dissociated from Arc-Cre/ER^T2^;Ai14 mice (Fig. 4A). According to a previous study (*26*), Gpr39, Aig1, Grin1, and Gpr137-expressing neurons in the BLA are involved in processing the positive valence, such as exposure to female mice (*26*). These genes were found in the saccharin-responsive neurons more than H_2_O-responsive neurons (Fig. 4B). Other positive valence-related genes, such as Ppp1r1b, Neurl1a, Esrra, and Thy1, expressed in both H_2_O- and saccharin-responsive neurons (Fig. 4B). Rspo2, Htr2c, Gabrg1, and Gabra1-expressing BLA neurons are reported to encode negative valence caused by an electrical foot shock (*26*). In agreement with this finding, expression of these genes were detected mainly in H_2_O-responsive neurons (Fig. 4C). Nptx2, Zfpm2, Acvr1c, and Gabra2, which are also expressed in the foot shock-activated BLA neurons (*26*), were not dominantly expressed in the H_2_O-responsive neurons (Fig. 4C). We also found each stimulus-specific gene; Fig. 4D and E show the top 10 of the lowest *p*-value genes comparing saccharin vs H_2_O. Saccharin-responsive neurons expressed ion channel, Cacna1f; Ras-related GTP-binding protein, Rragd; glycogen synthesis, Gys1; glutamate receptor, Grm8; lipid synthesis, Gpam (Fig. 4D). H_2_O-responsive neurons expressed cytoskeletal genes, Tmsb10 and Myh1/2; ribosomal gene, Rps15a; mitochondrial gene, Ndufs6; secretory gene, CCK (Fig. 4E). Serotonin plays a key role in modulating circuits involved in the regulation of emotion (*28*). Thus, we examined the expression of serotonin receptors, and detected the expression of Htr1f and Htr2b genes in saccharin-responsive neurons, along with Htr2c and Htr7 genes in H_2_O-responsive neurons (Fig. 4F). Taken together, our sequencing results suggest that genetically distinct clusters of BLA neurons respond to saccharin and H_2_O.

**Fig. 4.**
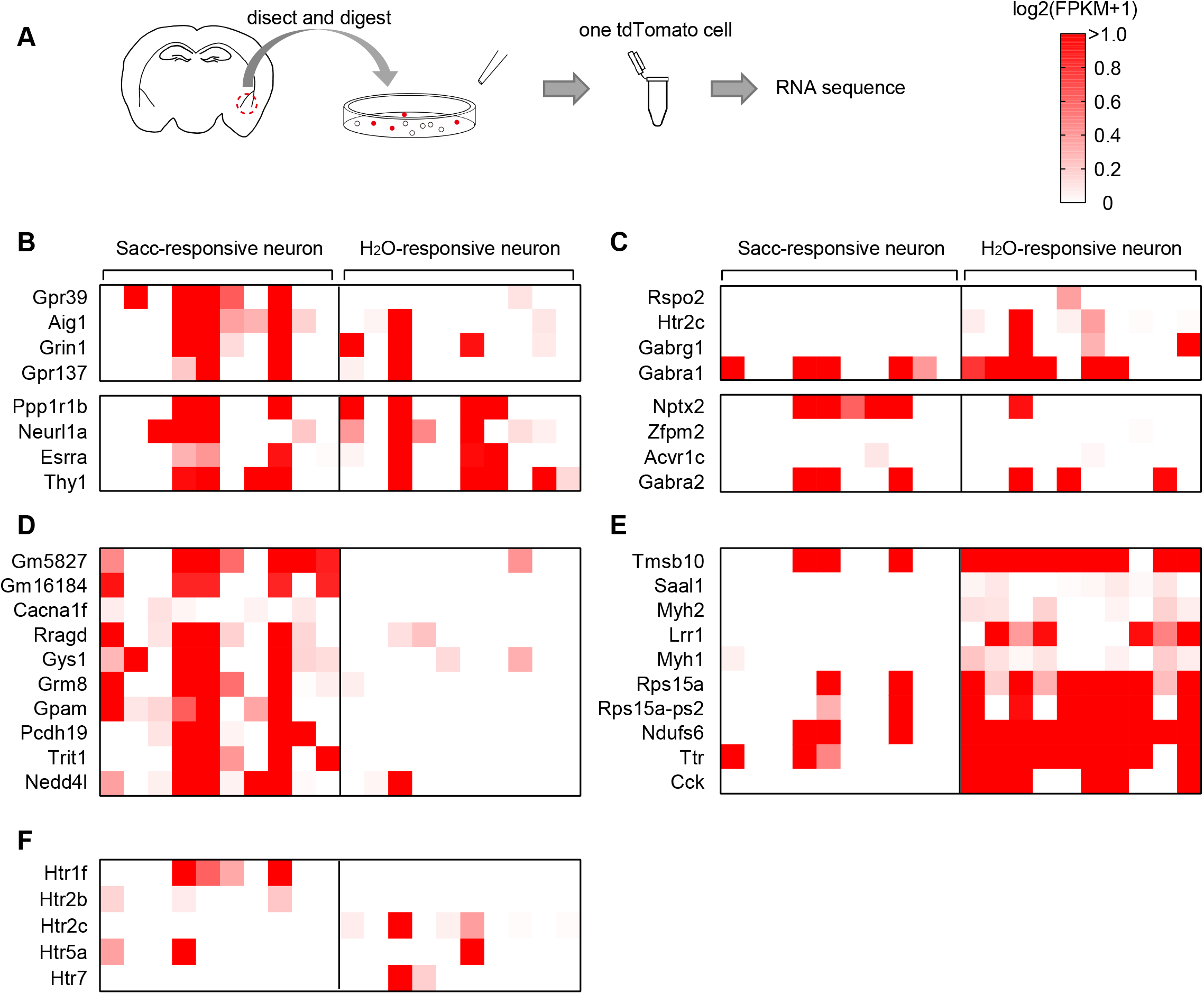
RNA sequence of TRAPed neurons in the BLA. (A) Experimental scheme of the RNA sequencing protocol. (B-C) Heatmap of the gene expression related to negative (B) and positive valence (C). Gene list was obtained from a previous report (*26*). Data are shown as Log2 (FPKM + 1). (D-E) H_2_O- (D) and saccharin-responsive neuron (E) specific gene expressions. (F) Heatmap of the gene expression of serotonin receptors.

## Discussion

Prediction of sugar intake is a key physiological response to preparing an efficient glucose utilization from the bloodstream (*29*). Here, we show that sucrose-conditioning and perception of sweet taste by saccharin are both necessary to increase insulin sensitivity. However, the unexpected zero-sweetness by water acutely inhibited insulin secretion without changing insulin sensitivity only when mice are conditioned with sucrose. These data suggest that the gap between expected- and actual sweetness is compared in the brain to modulate peripheral glucose metabolism. Cephalic phase insulin release (CPIR), a slight increase in insulin concentration before an increase in blood glucose after a meal, has been thought to be an important event to control postprandial glucose metabolism (*30*). Electrical lesion of the CeA and ventromedial nucleus of the hypothalamus abolishes CPIR (*31*, *32*). CPIR is mediated by preganglionic parasympathetic neurons within the dorsal motor nucleus of the vagus (*31*, *33*). BLA neurons have connections with CeA and VMH (*34*). These neuronal pathways may control peripheral glucose metabolism. However, the importance of CPIR on the regulation of blood glucose levels is still in debate (*35*). Thus, we focused on blood glucose level itself instead of CPIR.

We also found that these cephalic responses are mediated by distinct BLA neurons that have different gene expression patterns. Consistent with this, recent evidence indicates that distinct populations of BLA neurons, which have specific neuronal projections, encode positive and negative valences (*26*). Although many BLA research studies use electrical foot shock, quinine, or lithium chloride solutions to study negative valences and avoidance behavior, water-induced unmet-expectation was enough to change glucose metabolism without affecting the HPA axis, which represents strong psychological stress that can impair peripheral insulin sensitivity (*36*). Gene expressions in the saccharin- and H_2_O-responsive neurons partly, but not completely, overlap with the neurons that encode female- and foot shock-induced positive and negative valence respectively, suggesting that neurons encoding taste-eliciting valences share female-related and fear-related neuronal circuitry. Food perception is also achieved by olfactory and visual cues and thus, the cephalic response occurs when caged food is presented to the mice. Caged food rapidly activates neurons that have a pivotal role in appetite regulation, such as POMC neurons (*37*). Activation of POMC neurons increases sympathetic activity in the liver and changes hepatic lipid synthesis with modulation of morphological remodeling in the endoplasmic reticulum (*38*). The whole neuronal mechanism of the cephalic phase response remains to be elucidated.

Taken together, our data identified neurons in the BLA that are responsible for the systemic glucose metabolism, which is regulated by experience and prediction. The memory-based regulation also affects uncontrolled glucose metabolism induced by unexpected change in taste. The neuronal mechanism may be important for the systemic regulation of nutrition in daily life.

## Acknowledgments

We thank Takako Usuda for providing the mouse illustration and Nur Farehan Asgar, PhD, for editing a draft of this manuscript.

## Funding

Leading Initiative for Excellent Young Researchers (CT), Japan Agency for Medical Research and Development (AMED-RPIME, JP21gm6510009h0001) (CT), Grant-in-Aid for Young Scientists (A) JP17H05059 (CT), Grant-in-Aid for Scientific Research (B) JP21H02352 (CT), Grant-in-Aid for Scientific Research (B) JP18H02857 (NI), Grant-in-Aid for Challenging Research (Pioneering) 21K18275 (NI), the Takeda Science Foundation (CT), The Uehara Memorial Foundation (CT), Astellas Foundation for Research on Metabolic Disorders (CT), Suzuken Memorial Foundation (CT), Narishige Neuroscience Research Foundation (CT), Akiyama Life Science Foundation (CT), Program for supporting introduction of the new sharing system (JPMXS0420100617, JPMXS0420100618, JPMXS0420100619) (CT), Japanese Initiative for Progress of Research on Infectious Diseases for Global Epidemics (JP17fm0208011h0001, JP18fm0208011h0002, JP19fm0208011h0003) (CT)

## Author contributions

Conceptualization: IY, CT, Methodology: IY, YT, CT, Investigation: IY, TY, TI, SCX, CT, Visualization: IY, TY, Funding acquisition: NI, CT, Project administration: CT, Supervision: NI, KK, SD, Writing – original draft: IY, Writing – review & editing: SD, CT

## Notes

### Competing Interest Statement

The authors have declared no competing interest.

